# Progress Towards Plant Community Transcriptomics: Pilot RNA-Seq Data from 24 Species of Vascular Plants at Harvard Forest

**DOI:** 10.1101/2020.03.31.018945

**Authors:** Hannah E. Marx, Stacy A. Jorgensen, Eldridge Wisely, Zheng Li, Katrina M. Dlugosch, Michael S. Barker

## Abstract

- *Premise of the study:* Large scale projects such as NEON are collecting ecological data on entire biomes to track and understand plant responses to climate change. NEON provides an opportunity for researchers to launch community transcriptomic projects that ask integrative questions in ecology and evolution. We conducted a pilot study to investigate the challenges of collecting RNA-seq data from phylogenetically diverse NEON plant communities, including species with diploid and polyploid genomes.
- *Methods:* We used Illumina NextSeq to generate >20 Gb of RNA-seq for each of 24 vascular plant species representing 12 genera and 9 families at the Harvard Forest NEON site. Each species was sampled twice, in July and August 2016. We used Transrate, BUSCO, and GO analyses to assess transcriptome quality and content.
- *Results:* We obtained nearly 650 Gb of RNA-seq data that assembled into more than 755,000 translated protein sequences across the 24 species. We observed only modest differences in assembly quality scores across a range of k-mer values. On average, transcriptomes contained hits to >70% of loci in the BUSCO database. We found no significant difference in the number of assembled and annotated genes between diploid and polyploid transcriptomes.
- *Discussion:* Our resource provides new RNA-seq datasets for 24 species of vascular plants in Harvard Forest. Challenges associated with this type of study included recovery of high quality RNA from diverse species and access to NEON sites for genomic sampling. Overcoming these challenges offers clear opportunities for large scale studies at the intersection of ecology and genomics.

## INTRODUCTION

Many questions in ecology and evolutionary biology increasingly require combining data from these fields at large scales. In particular, integrated, large-scale analyses of multispecies ecological and phylogenetic data sets have become critical to understanding plant distributions and responses to climate change (Zanne et al., 2014; Swenson and Jones, 2017; Maitner et al., 2018; Enquist et al., 2019; Gallagher et al., 2019; McFadden et al., 2019; Rice et al., 2019; Baniaga et al., 2020; Román-Palacios and Wiens, 2020). Recognizing this need, NSF recently launched the National Ecological Observatory Network (NEON) to generate large-scale data on species occurrence, phenology, climate, and more, for ecological communities across the United States (Collinge, 2018; Knapp and Collins, 2019). Metagenomic and genomic sampling is also being used to identify and estimate changes in abundance and composition of some taxa, especially microbial communities (https://www.neonscience.org/data). Although these data and analyses will be crucial for understanding ecosystem scale processes, collection of genomic data from a broader array of species across NEON sites would allow researchers to further integrate ecological and evolutionary processes in the analyses of communities.

Genomic analyses of single species, although important, do not capture the larger patterns occurring within an interacting community of plants. Trancriptome profiling or genome sequencing of multiple species and individuals within a community will open new, integrative avenues of analyses and allow us to address existing questions that require sampling of floras and communities (Bragg et al., 2015; Fitzpatrick and Keller, 2015; Bowsher et al., 2017; Han et al., 2017; Swenson and Jones, 2017; Zambrano et al., 2017; Matthews et al., 2018; Subrahmaniam et al., 2018; Breed et al., 2019). This is especially true for understanding responses to climate change where community level analyses are needed to capture the interacting dynamics of different species responses (Liu et al., 2018; Komatsu et al., 2019; Snell et al., 2019). The integration of community level genomic data from non-model species with ecological and trait data will improve our understanding of plant responses to climate change. Collecting genomic data at the community level with repeated sampling that mirrors other trait data collection will permit assessments of the genetic diversity of entire plant communities and how they change over time, estimates of gene flow and hybridization, measurement of *in situ* gene expression variation across species in response to shared climate events, and a genomic perspective on functional diversity within and between plant communities. Metagenomics analyses of microbiomes have transformed our understanding of and approaches for studying microbial biology (Fierer, Lauber, et al., 2012; Fierer, Leff, et al., 2012; Turner et al., 2013; Delgado-Baquerizo et al., 2018; Jansson and Hofmockel, 2020). Similar plant community transcriptomics and genomics studies could open new avenues of research and provide the crucial data to understand plant responses to climate change.

To explore the potential and challenges of plant community transcriptomics, we conducted a pilot RNA-seq study at the Harvard Forest NEON site (HARV). We sampled 24 species of vascular plants from sites adjacent to the NEON plot. Species were selected from a phylogenetically diverse range of plants that included ferns, trees, and herbaceous annuals. For plants in particular, the abundance of polyploid species in communities has the potential to pose challenges for genomic studies. To explore the impacts of polyploidy on transcriptome surveys, we made an effort to select sets of related polyploid and diploid species. Each species was sampled at two different time points in July and August 2016. Here, we give an overview of our data collection, present new reference transcriptomes and translated protein collections for each species, and evaluate the quality of these assemblies using multiple approaches.

## METHODS

### Taxon selection and sampling

The Harvard Forest Flora (Jenkins et al., 2008) was used to select taxa to represent each category (native/invasive, diploid/polyploid). Invasive species status was determined from the Harvard Forest Flora Database (Jenkins and Motzkin, 2009). Putative diploids and neo-polyploid species were identified from chromosome counts obtained from the Chromosome Counts Database (Rice et al., 2015). Congeneric species pairs were selected based on their phylogenetic relatedness. Our sampling included nine polyploid and eleven diploid species (Table 1). We could not determine the ploidal level of four species. The Harvard Forest Flora Database was used to locate sampling sites.

**Table 1.** Summary statistics for RNA-seq data sets, assemblies with a k-mer = 57, and translations. P and D are polyploid and diploid species, respectively.

| Species | Ploidy | Chrom.<br># | July SRA | Aug SRA | Total Gb<br>(July +<br>Aug) | # of<br>Scaffolds | Mean<br>Scaffold<br>Length<br>(bp) | Scaffold<br>N50 (bp) | # of<br>Translated<br>Proteins | Mean<br>Translated<br>Length<br>(nucleic<br>acids, bp) | N50<br>(trans.<br>nucleic<br>acids, bp) |
| --- | --- | --- | --- | --- | --- | --- | --- | --- | --- | --- | --- |
| *Dryopteris carthusiana* | P | 82 | SAMN0827<br>7176 | SAMN0827<br>7204 | 26 | 643129 | 264.5 | 710 | 24851 | 473.8 | 543 |
| *Dryopteris intermedia* | D | 41 | SAMN0827<br>7187 | SAMN0827<br>7216 | 22 | 529510 | 267.1 | 822 | 22595 | 587.1 | 702 |
| *Dryopteris marginalis* | D | 41 | SAMN0827<br>7188 | SAMN0827<br>7217 | 21 | 550548 | 260.1 | 917 | 34121 | 633.9 | 789 |
| *Galium mollugo* | D | 11 | SAMN0827<br>7173 | SAMN0827<br>7201 | 40 | 608764 | 179.2 | 1028 | 25040 | 692.9 | 822 |
| *Galium tinctorium* | D | 12 | SAMN0827<br>7181 | SAMN0827<br>7210 | 32 | 78487 | 400.6 | 1091 | 16610 | 811.2 | 1023 |
| *Galium triflorum* | P | 33 | SAMN0827<br>7189 | SAMN0827<br>7219 | 37 | 574562 | 246 | 899 | 38650 | 664.9 | 798 |
| *Hypericum perforatum* | P | 16 | SAMN0827<br>7171 | SAMN0827<br>7199 | 25 | 335837 | 233.4 | 867 | 34670 | 698.6 | 858 |
| *Juglans cinerea* | D | 16 | SAMN0827<br>7174 | SAMN0827<br>7202 | 18.2 | 569859 | 359.5 | 1151 | 66595 | 712.4 | 918 |
| *Lonicera tatarica*<br>*var. morrowii* | NA | 9 | SAMN0827<br>7167 | SAMN0827<br>7195 | 10.4 | 386927 | 324.9 | 1216 | 28147 | 710.9 | 864 |
| *Lysimachia ciliata* | P | 48 | SAMN0827<br>7190 | SAMN0827<br>7220 | 24 | 1422451 | 207 | 828 | 32005 | 631.6 | 777 |
| *Lysimachia nummularia* | D | 17 | SAMN0827<br>7194 | SAMN0827<br>7223 | 25.3 | 428232 | 320.9 | 1102 | 46343 | 716.7 | 894 |
| *Lysimachia quadrifolia* | P | 42 | SAMN0827<br>7183 | SAMN0827<br>7212 | 30 | 340491 | 290.4 | 944 | 37737 | 674.9 | 813 |
| *Persicaria arifolia* | NA | NA | SAMN0827<br>7180 | SAMN0827<br>7209 | 30 | 528292 | 245 | 787 | 28741 | 558.8 | 639 |
| *Persicaria hydropiperoides* | NA | NA | SAMN0827<br>7172 | SAMN0827<br>7200 | 21.3 | 347558 | 344 | 913 | 46058 | 658.8 | 795 |
| *Persicaria sagittata* | NA | 20 | SAMN0827<br>7179 | SAMN0827<br>7208 | 26 | 398304 | 319.6 | 1148 | 33118 | 725.1 | 885 |
| *Plantago lanceolata* | D | 6 | SAMN0827<br>7178 | SAMN0827<br>7206 | 31 | 213834 | 293.1 | 1073 | 20470 | 742.7 | 906 |
| *Plantago major* | D | 6 | SAMN0827<br>7169 | SAMN0827<br>7197 | 23 | 217041 | 308.6 | 1032 | 24715 | 782.6 | 993 |
| *Plantago rugelii* | P | 12 | SAMN0827<br>7170 | SAMN0827<br>7198 | 30 | 378418 | 276.1 | 1161 | 30673 | 760.5 | 930 |
| *Polygonum cilinode* | D | 11 | SAMN0827<br>7184 | SAMN0827<br>7213 | 17.3 | 1065186 | 186.8 | 415 | 6088 | 471.6 | 498 |
| *Potentilla argentea* | P | 21 | SAMN0827<br>7177 | SAMN0827<br>7207 | 29 | 245734 | 425.6 | 1268 | 16306 | 525.5 | 687 |
| *Potentilla canadensis* | D | 14 | SAMN0827<br>7192 | SAMN0827<br>7222 | 16.6 | 433249 | 216.5 | 667 | 37503 | 526.5 | 594 |
| *Prunus serotina* | P | 16 | SAMN0827<br>7186 | SAMN0827<br>7215 | 31 | 350572 | 267.5 | 1017 | 30812 | 658.3 | 774 |
| *Prunus virginiana* | D | 8 | SAMN0827<br>7185 | SAMN0827<br>7214 | 38 | 536216 | 235 | 1110 | 38773 | 678.8 | 813 |
| *Reynoutria japonica* | P | 33 | SAMN0827<br>7193 | SAMN0827<br>7224 | 44 | 410810 | 279.6 | 870 | 34662 | 549.3 | 609 |

Tissue from mature leaves was collected from an individual representing each target species at two time points (July and August) during the 2016 growing season. The same individual was sampled at both time points for perennial individuals, and the same population was sampled for annuals. Field sampling for plant RNA-seq followed the protocol described in Yang *et al*. 2017 (Yang et al., 2017). Leaf tissues were flash frozen in liquid nitrogen in the field, and shipped on dry ice to the University of Arizona for RNA extraction.

### RNA extraction and RNA-seq

Total RNA was extracted from leaf tissue collected at each time point for all species using the Spectrum Plant Total RNA Kit (Sigma-Aldrich Co., St. Louis, MO, USA) following Protocol A. RNA was used to prepare cDNA using Nugen’s Ovation RNA-Seq System via single primer isothermal amplification (Catalogue # 7102-A01) and automated on the Apollo 324 liquid handler (Wafergen). cDNA was quantified on the Nanodrop (Thermo Fisher Scientific) and was sheared to approximately 300 bp fragments using the Covaris M220 ultrasonicator. Libraries were generated using Kapa Biosystem’s library preparation kit (KK8201). Fragments were end repaired and A-tailed, and individual indexes and adapters (Bioo, catalogue #520999) were ligated on each separate sample. The adapter ligated molecules were cleaned using AMPure beads (Agencourt Bioscience/Beckman Coulter, A63883), and amplified with Kapa’s HIFI enzyme (KK2502). Each library was then analyzed for fragment size on an Agilent’s Tapestation, and quantified by qPCR (KAPA Library Quantification Kit, KK4835) on Thermo Fisher Scientific’s Quantstudio 5 before multiplex pooling (13-16 samples per lane) and paired-end sequencing at 2×150 bp on the Illumina NextSeq500 platform at Arizona State University’s CLAS Genomics Core facility. Raw read quality was assessed using fastQC (Andrews, 2010).

### *De novo* transcriptome assembly, protein translation, and quality assessment

Raw sequence reads were processed using the SnoWhite pipeline (Barker et al., 2010; Dlugosch et al., 2013), which included trimming adapter sequences and bases with a quality score below 20 from the 3’ ends of all reads, removing reads that are entirely primer and/or adapter fragments using TagDust (Lassmann et al., 2009), and removing polyA/T tails with SeqClean (https://sourceforge.net/projects/seqclean/). The cleaned reads from each sample time point were merged and cleaned to synchronize read pairs using fastq-pair (Edwards and Edwards, 2019), and pooled to assemble a reference *de novo* transcriptome for each species.

Due to the significant time involved in running and evaluating multiple assemblies for each species, we chose five species (*Dryopteris intermedia, Galium mollugo, Juglans cinerea, Plantago major*, and *Persicaria sagittata*) to identify the optimal k-mer to use for assembling all 24 species. For these five exemplar taxa, we examined the quality of assemblies generated by SOAPdenovo-Trans v1.03 (Xie et al., 2014) across a range of k-mers (37, 47, 57, 67, 77, 87, 97, 107, 117, and 127). Assembly quality across the different k-mers was assessed by mapping the raw reads to each assembly with Transrate v1.0.3 (Smith-Unna et al., 2016) and evaluating the optimal assembly scores. Transrate calculates assembly scores by remapping the reads back to the assembly and combining a variety of metrics for each contig including estimates of whether a base pair was called correctly, whether a base should be a part of the final transcript, the probability that a contig was derived from a single transcript, and the probability that a contig is structural complete. We selected a k-mer that produced the average highest optimal assembly score across the five species. This k-mer (57, see Results) was used to assemble reference transcriptomes for the entire collection of species.

We used TransPipe (Barker et al., 2010) to identify plant proteins within the assembled transcripts for each reference transcriptome and provide protein and in-frame nucleic acid sequences for each species. The reading frame and protein translation for each sequence was identified by comparison to protein sequences from 25 sequenced and annotated plant genomes from Phytozome (Goodstein et al., 2012). Using BLASTX (Wheeler et al., 2008), best hit proteins were paired with each gene at a minimum cutoff of 30% sequence similarity over at least 150 sites. Genes that did not have a best hit protein at this level were removed. To determine the reading frame and generate estimated amino acid sequences, each gene was aligned against its best hit protein by Genewise 2.2.2 (Birney et al., 2004). Based on the highest scoring Genewise DNA-protein alignments, stop and ‘N’ containing codons were removed to produce estimated amino acid sequences for each gene. Output included paired DNA and protein sequences with the DNA sequence reading frame corresponding to each protein sequence.

To assess the quality of the assembled transcriptomes for the full set of 24 species, we analyzed each with Transrate and BUSCO. Summary statistics including the number of scaffolds, mean scaffold lengths, and N50 were calculated by Transrate v1.0.3 for all scaffolds as well as the subset of sequences that were identified as plant proteins and translated. We evaluated the completeness of our transcriptome coverage with BUSCO v4.0.5 (Seppey et al., 2019). BUSCO compares sequences to a collection of universal single copy orthologs for the viridiplantae (Viridiplantae Odb10) and the eukaryotes (Eukaryote Odb10). We also used the Transrate and BUSCO statistics to compare differences in the assemblies of diploid and polyploid species.

### Gene Ontology (GO) Annotation and Comparison

Gene Ontology annotations of all transcriptomes were obtained through Translated BLAST (Blastx) searches against annotated *A. thaliana* protein database from TAIR (Lamesch et al., 2012) to find the best hit with length of at least 100 bp and an e*-*value of at least 1e-10. GO annotations were obtained for the whole transcriptome for each species and presented as a heatmap. The heatmap columns were clustered by hierarchical clustering with default parameters in R. The overall ranking of GO category rows was determined by the ranking of GO annotations among the transcriptome of *Lysimachia ciliata*, as an arbitrary standard. The colors of the heatmap represent the percentage of the transcriptome represented by a particular GO category, with red being highest and purple lowest.

## RESULTS

We found relatively little variation in the optimal Transrate scores across assemblies with different k-mers. The optimal Transrate scores ranged from ∼0.1–0.15 with each of the five exemplar species peaking at different k-mers (Figure 1). Scores trended downward for all species at higher k-mers with no sharp peaks in the score apparent in most taxa. The mean k-mer of the top scoring assemblies for each species was 61, and the closest k-mer to this value (57) was used to assemble reference transcriptomes for all 24 species.

**Figure 1.**
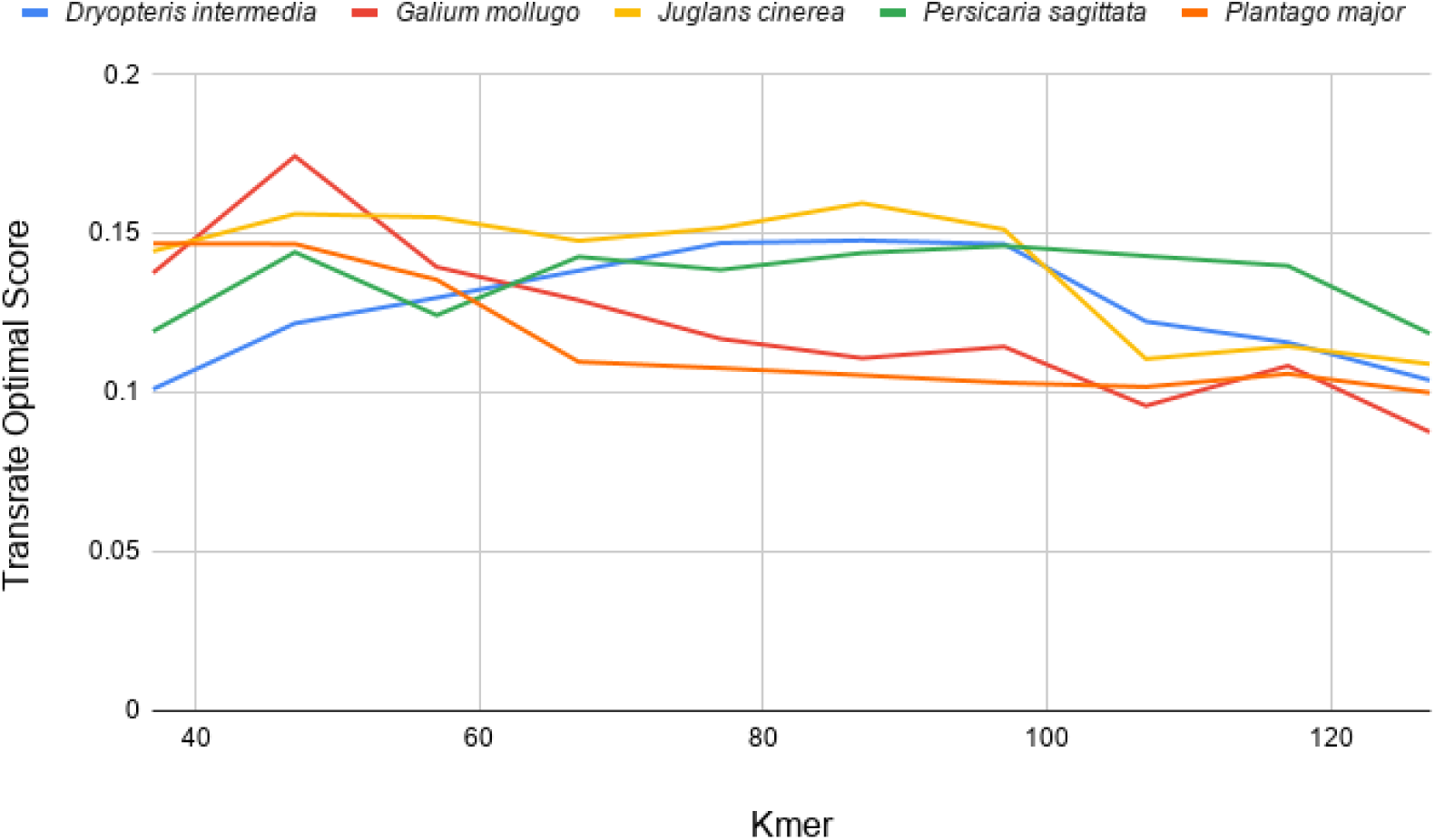
Transrate Optimal Scores for assemblies of five exemplar species from Harvard Forest. A reference transcriptome for each species was assembled with different K-mers starting at K = 37 and increasing in increments of 10 to K = 127.

Assemblies for most of the 24 species appeared to be relatively high quality. We achieved a mean read depth of 27 Gb for each reference transcriptome (Table 1). With a k-mer = 57, the assemblies contained an average of 483,084 scaffolds with a mean length of 281 bp and N50 of 960 bp. The translated nucleic acids for each assembly had an average of 31,470 sequences with a mean length of 652 bp and N50 of 789 bp. We observed no significant relationship between the number of scaffolds or number of translated proteins and sequencing depth (Figure 2), suggesting that our sequencing effort on these libraries was sufficient. The mean complete + fragmented BUSCO percentages were 73.2% against the viridiplantae database and 76% against the eukaryote database (Table 2). We found more hits to sequences in the BUSCO databases with more translated proteins, but the hits plateaued around 20,000 proteins (Figure 3).

**Table 2.** BUSCO results for comparisons to the viridiplantae and eukaryote databases. C = % all complete, S = % complete and single copy, D = % complete and duplicated, F = % fragmented, M = % missing, D/S = ratio of duplicated to single copy complete sequences, and C+F = % of complete and fragmented BUSCO matches in the respective database.

|  | BUSCO Viridiplantae DB |  |  |  |  |  |  | BUSCO Eukaryote DB |  |  |  |  |  |  |
| --- | --- | --- | --- | --- | --- | --- | --- | --- | --- | --- | --- | --- | --- | --- |
| Species | C | (S) | (D) | F | M | D/S | C+F | C | (S) | (D) | F | M | D/S | C+F |
| *Dryopteris carthusiana* | 19.3 | 16.9 | 2.4 | 40.7 | 40 | 0.142 | 60 | 24.3 | 20 | 4.3 | 43.5 | 32.2 | 0.215 | 67.8 |
| *Dryopteris intermedia* | 39.8 | 35.1 | 4.7 | 38.4 | 21.8 | 0.134 | 78.2 | 48.2 | 44.3 | 3.9 | 32.9 | 18.9 | 0.088 | 81.1 |
| *Dryopteris marginalis* | 41.5 | 34.4 | 7.1 | 40 | 18.5 | 0.206 | 81.5 | 48.6 | 43.5 | 5.1 | 38.4 | 13 | 0.117 | 87 |
| *Galium mollugo* | 27.8 | 22.6 | 5.2 | 50.1 | 22.1 | 0.230 | 77.9 | 38 | 32.5 | 5.5 | 41.6 | 20.4 | 0.169 | 79.6 |
| *Galium tinctorium* | 40 | 37.9 | 2.1 | 40 | 20 | 0.055 | 80 | 50.2 | 46.3 | 3.9 | 27.1 | 22.7 | 0.084 | 77.3 |
| *Galium triflorum* | 30.8 | 21.9 | 8.9 | 46.4 | 22.8 | 0.406 | 77.2 | 33.4 | 21.6 | 11.8 | 50.2 | 16.4 | 0.546 | 83.6 |
| *Hypericum perforatum* | 29.5 | 22.4 | 7.1 | 48.7 | 21.8 | 0.317 | 78.2 | 38.8 | 29 | 9.8 | 40 | 21.2 | 0.338 | 78.8 |
| *Juglans cinerea* | 33.9 | 27.8 | 6.1 | 48.2 | 17.9 | 0.219 | 82.1 | 47.8 | 39.2 | 8.6 | 34.1 | 18.1 | 0.219 | 81.9 |
| *Lonicera tatarica* var. *morrowii* | 34.8 | 29.4 | 5.4 | 46.4 | 18.8 | 0.184 | 81.2 | 47.1 | 36.9 | 10.2 | 34.5 | 18.4 | 0.276 | 81.6 |
| *Lysimachia ciliata* | 34.1 | 28.7 | 5.4 | 46.1 | 19.8 | 0.188 | 80.2 | 49 | 40 | 9 | 32.2 | 18.8 | 0.225 | 81.2 |
| *Lysimachia nummularia* | 42.4 | 35.1 | 7.3 | 42.8 | 14.8 | 0.208 | 85.2 | 48.6 | 34.9 | 13.7 | 37.6 | 13.8 | 0.393 | 86.2 |
| *Lysimachia quadrifolia* | 31.8 | 26.4 | 5.4 | 44.2 | 24 | 0.205 | 76 | 38 | 32.5 | 5.5 | 39.6 | 22.4 | 0.169 | 77.6 |
| *Persicaria arifolia* | 13.9 | 10.6 | 3.3 | 51.5 | 34.6 | 0.311 | 65.4 | 24.7 | 16.9 | 7.8 | 46.3 | 29 | 0.462 | 71 |
| *Persicaria hydropiperoides* | 25.4 | 16.5 | 8.9 | 48.9 | 25.7 | 0.539 | 74.3 | 32.9 | 18.4 | 14.5 | 44.3 | 22.8 | 0.788 | 77.2 |
| *Persicaria sagittata* | 27.8 | 16.5 | 11.3 | 48.7 | 23.5 | 0.685 | 76.5 | 39.2 | 20.4 | 18.8 | 40.8 | 20 | 0.922 | 80 |
| *Plantago lanceolata* | 40.5 | 36 | 4.5 | 40.5 | 19 | 0.125 | 81 | 45.9 | 39.2 | 6.7 | 36.1 | 18 | 0.171 | 82 |
| *Plantago major* | 51.3 | 48 | 3.3 | 33.4 | 15.3 | 0.069 | 84.7 | 54.1 | 49.8 | 4.3 | 29.4 | 16.5 | 0.086 | 83.5 |
| *Plantago rugelii* | 34.5 | 20.9 | 13.6 | 46.6 | 18.9 | 0.651 | 81.1 | 45.8 | 27.8 | 18 | 39.6 | 14.6 | 0.647 | 85.4 |
| *Polygonum cilinode* | 11.5 | 9.4 | 2.1 | 8.5 | 80 | 0.223 | 20 | 5.5 | 4.7 | 0.8 | 19.6 | 74.9 | 0.170 | 25.1 |
| *Potentilla argentea* | 11.7 | 10.8 | 0.9 | 33.2 | 55.1 | 0.083 | 44.9 | 14.1 | 12.9 | 1.2 | 38.4 | 47.5 | 0.093 | 52.5 |
| *Potentilla canadensis* | 12.9 | 9.6 | 3.3 | 47.8 | 39.3 | 0.344 | 60.7 | 17.3 | 12.2 | 5.1 | 52.9 | 29.8 | 0.418 | 70.2 |
| *Prunus serotina* | 23.5 | 18.1 | 5.4 | 50.4 | 26.1 | 0.298 | 73.9 | 28.6 | 20 | 8.6 | 47.1 | 24.3 | 0.430 | 75.7 |
| *Prunus virginiana* | 27 | 18.1 | 8.9 | 54.4 | 18.6 | 0.492 | 81.4 | 37.7 | 25.5 | 12.2 | 44.7 | 17.6 | 0.478 | 82.4 |
| *Reynoutria japonica* | 30.1 | 22.8 | 7.3 | 45.2 | 24.7 | 0.320 | 75.3 | 30.2 | 23.1 | 7.1 | 45.5 | 24.3 | 0.307 | 75.7 |

**Figure 2.**
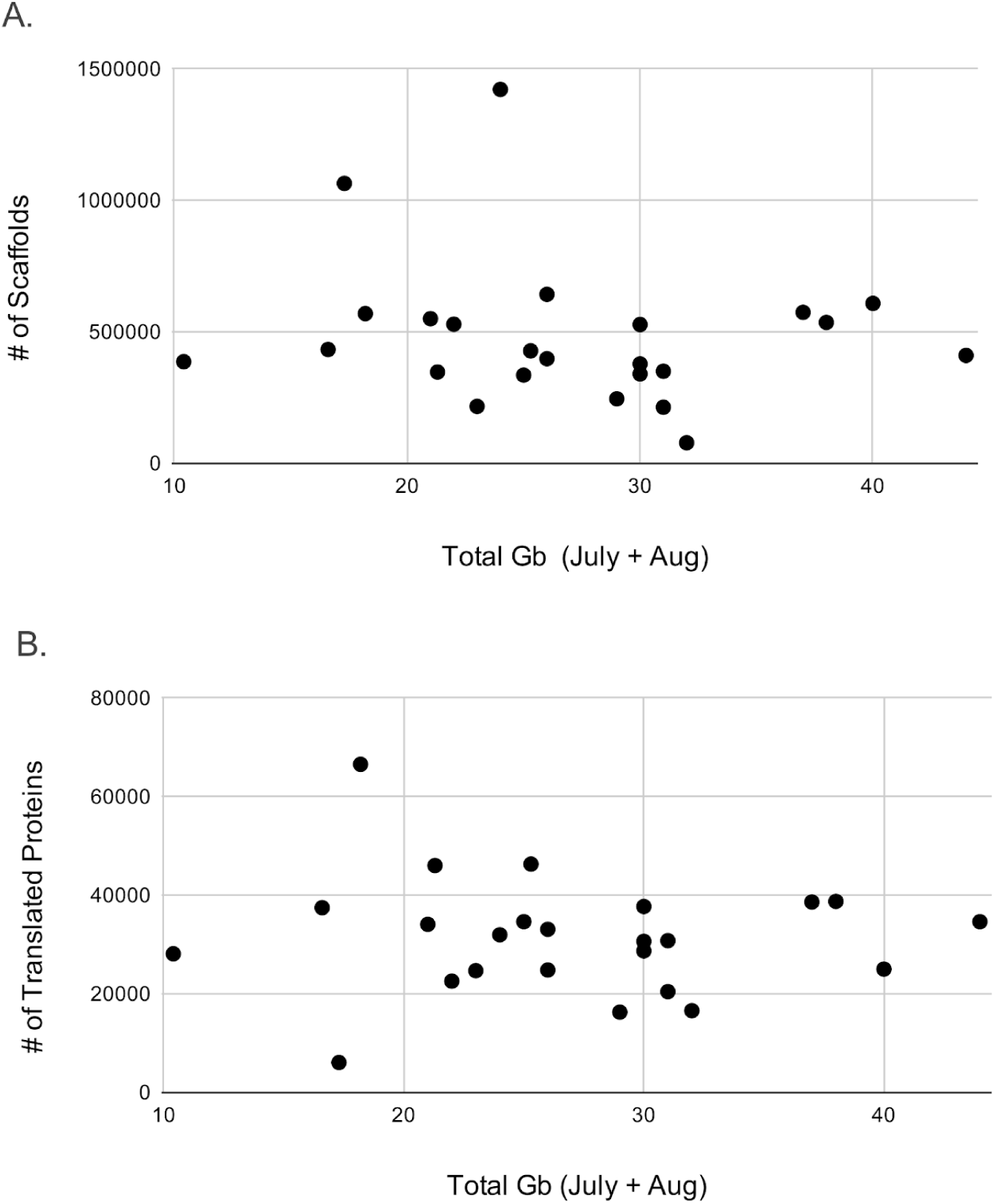
Comparison of the number of A) scaffolds and B) translated proteins produced by each assembly with the total gigabases sequenced for each species.

**Figure 3.**
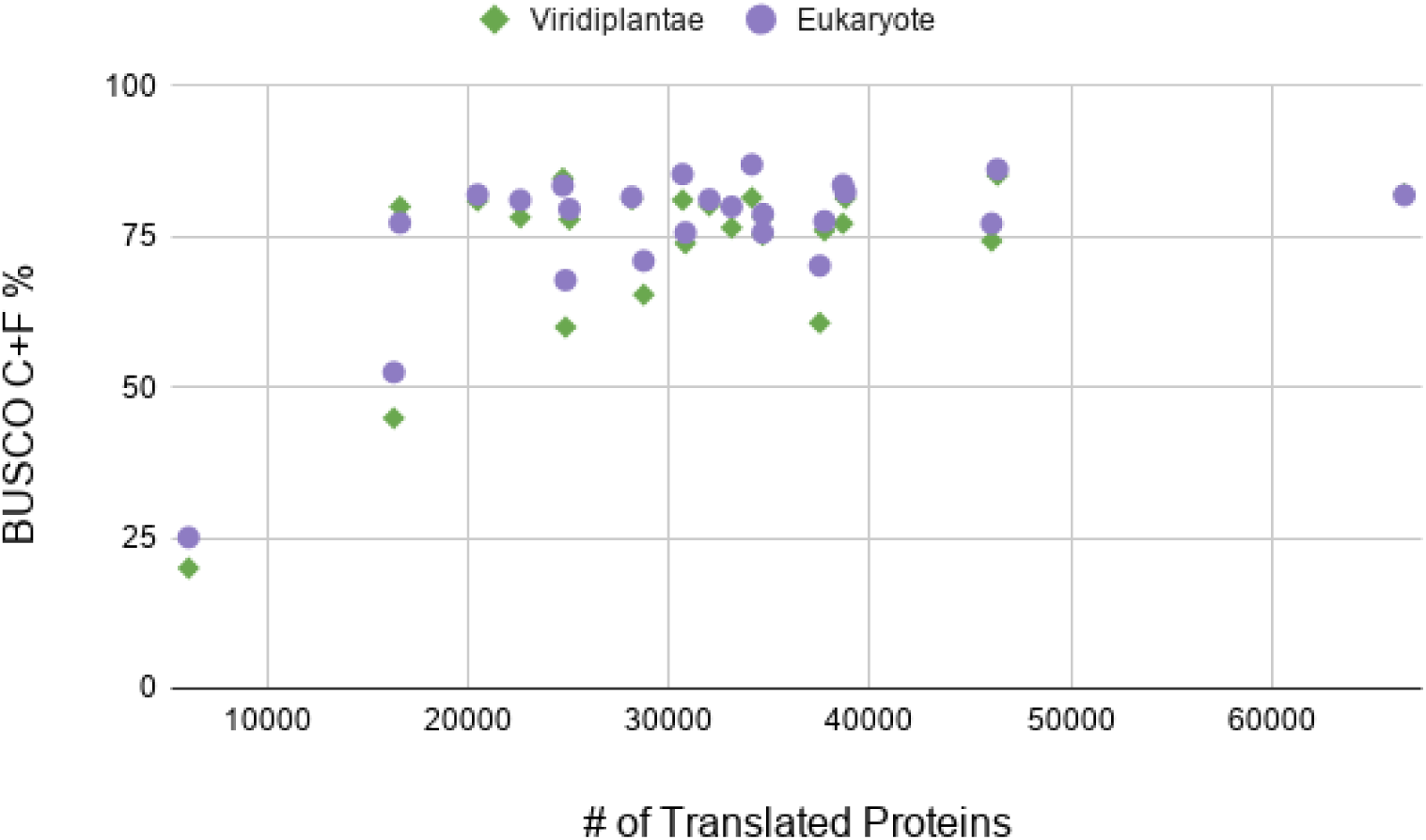
The percentage of BUSCO complete (C) plus fragmented (F) matches compared to the number of translated proteins in each assembly. Green diamonds represent BUSCO matches to the viridiplantae database, whereas purple circles represent matches to the eukaryote database.

Polyploid species did not have significantly more translated proteins than diploids with 31,152 average proteins translated compared to 30,804 (Figure 4A; two-tailed t-test *p* = 0.95). Similarly, polyploid species did not have a significantly higher proportion of duplicated BUSCO matches than diploids (Figure 4B; two-tailed t-test *p* = 0.11). In some cases the number of proteins or duplicated BUSCO proportion was lower when comparing a polyploid species with its related diploids (e.g., *Dryopteris*). This may be due to variation in read and/or assembly quality rather than differences in the biology of these species. However, it is not clear that this is due to differences in data quality because in most cases, including *Dryopteris*, all of the species have similarly high read depth (>20 Gb).

**Figure 4.**
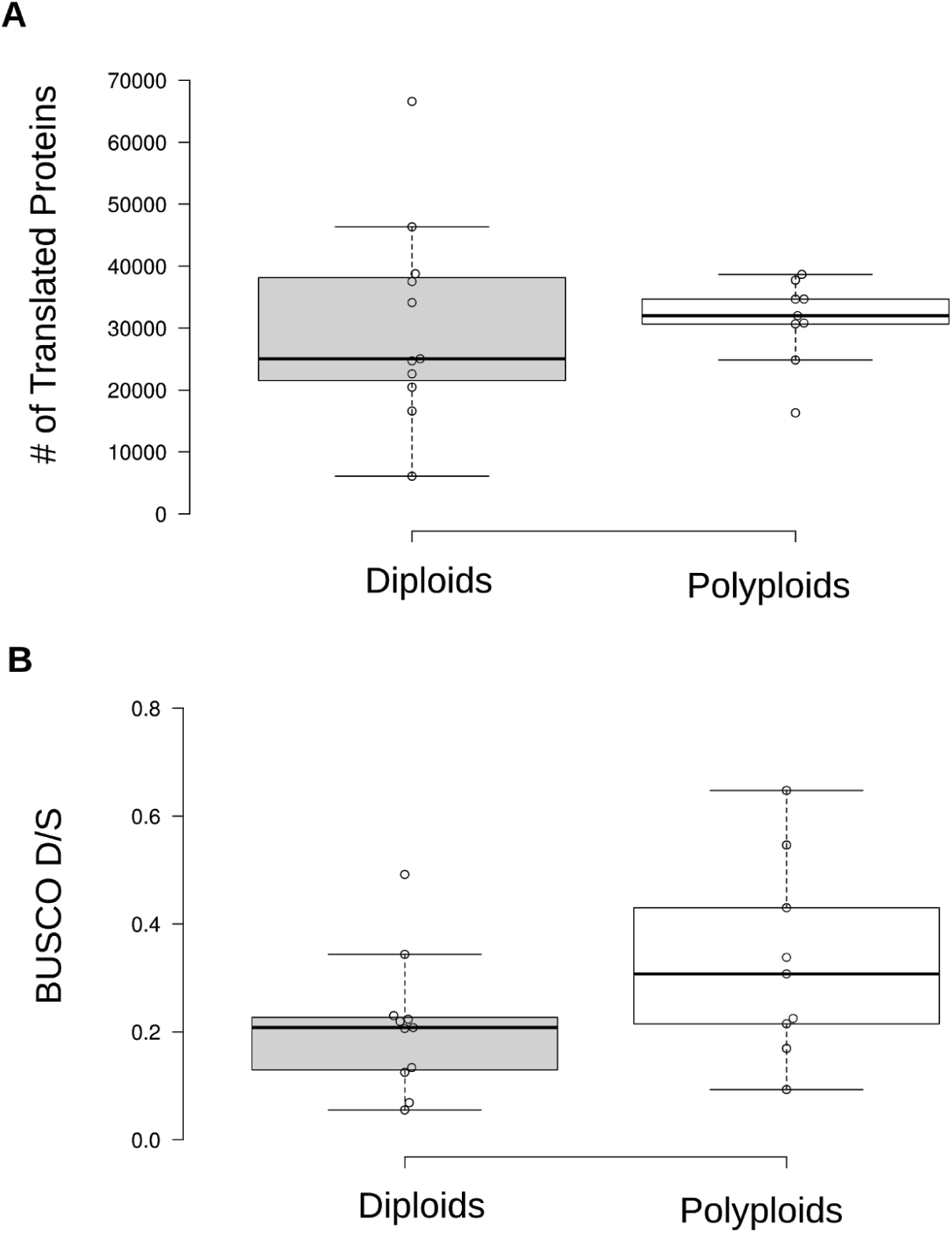
Comparison of A) the number of translated proteins and B) BUSCO duplicated/single copy ratio for assemblies of diploid and polyploid species. In neither case were diploids significantly different from polyploids.

Gene ontology (GO) annotations of the transcriptomes of the 24 species were largely similar (Figure 5). Categories such as “other cellular processes”, “other metabolic processes”, and “other intracellular components” were the largest fraction of all transcriptomes, whereas “receptor binding or activity” and “electron transport or energy pathways” were among the smallest. The rank order of each GO slim category was largely consistent across most species. Species from the same genus were sometimes clustered together, such as in *Dryopteris* and *Lysimachia*, but in most cases the species were not clustered with their congeners. *Polygonum cilinode* was unique in having many differences in GO category rank compared to the other taxa. It was also the lowest scoring transcriptome assembly with only 6,088 translated proteins and nearly 80% of BUSCO genes missing (Table 2).

**Figure 5.**
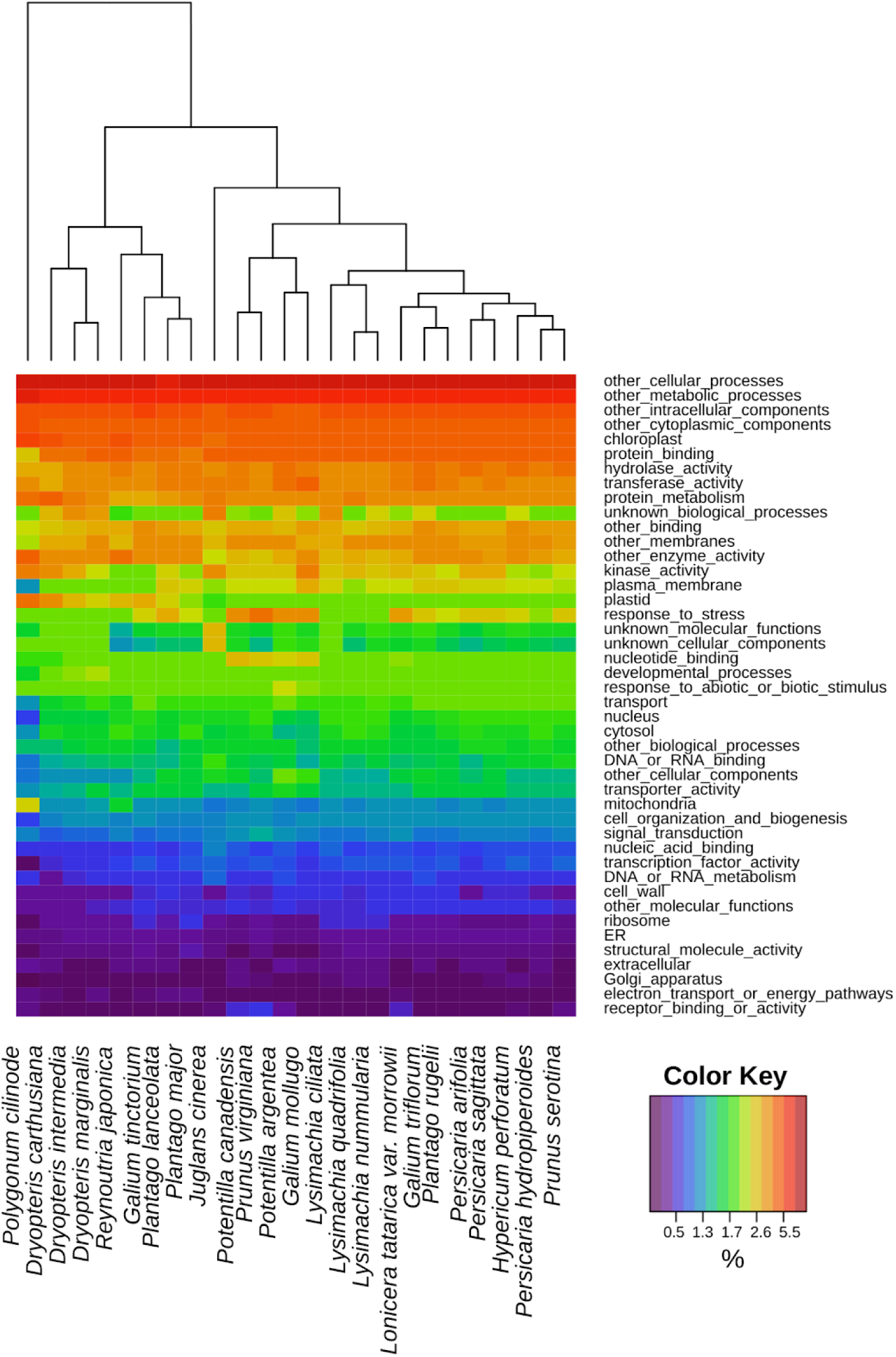
Heat map of gene ontology (GO) slim categories present in the entire transcriptome of each species. Each column represents the annotated GO categories from each analyzed transcriptome, whereas the rows represent a particular GO category. The colors of the heat map represent the percentage of the transcriptome represented by a particular GO category, with red being highest and purple lowest. The overall ranking of GO category rows was determined by the ranking of GO annotations in the transcriptome of *Lysimachia ciliata*. Hierarchical clustering was used to organize the heatmap columns.

## DISCUSSION

Overall, the transcriptomes we assembled for 24 species of vascular plants at Harvard Forest appear to be relatively high quality and consistent with our expectations for plant transcriptomes. In particular, the number of scaffolds that matched known plant proteins was consistent with the number of genes in sequenced plant genomes (Michael, 2014; Wendel et al., 2016). For example, our transcriptomes of *Prunus virginiana* and *P. serotina* contained 38,773 and 30,812 translated proteins each. Genomes of related *Prunus* species had similar numbers of annotated genes, including 27,852 in *P. persica* (International Peach Genome Initiative et al., 2013), *41,294 in P. yedoensis* (Baek et al., 2018), *and 43,349 in P. avium* (Shirasawa et al., 2017). The assemblies are also reasonably complete with more than 70% of BUSCO genes present on average. This is a similar distribution of BUSCO scores to those in the recently published 1KP project (Carpenter et al., 2019; One Thousand Plant Transcriptomes Initiative, 2019) and other studies (Blande et al., 2017; Evkaikina et al., 2017; Pokorn et al., 2017; Weisberg et al., 2017). Like many transcriptome assemblies (Johnson et al., 2012; Carpenter et al., 2019; Patterson et al., 2019), these assemblies also contain a large number of small scaffolds (<300 bp). Small scaffolds are likely artifacts of library amplification and sequencing, considering that most did not translate to a known plant protein sequence.

We found no significant difference in the number of translated genes between the diploid and polyploid transcriptome assemblies. Although this could be due to the modest sample size, it may also reflect biological differences in expressed transcriptome size and diversity that impact the number of assembled genes. Under a simple null model of polyploid transcriptome size, one may expect to observe an approximate doubling of the diploid transcriptome size that may translate to doubling the number of assembled genes. However, recent research indicates polyploid transcriptomes may be smaller than expected. Research in *Glycine* has found that the expressed transcriptome size of polyploid species are less than 2X the diploid size (Coate and Doyle, 2015; Doyle and Coate, 2018). For example, the transcriptome of the allotetraploid *Glycine dolichocarpa* was 1.4X the size of its diploid progenitors (Coate and Doyle, 2010). The apparent lower than expected level of the quantity of gene expression in polyploids may be an artifact of comparing diploids and polyploids without accounting for differences in cell numbers or biomass (Visger et al., 2019). However, smaller transcriptome sizes in polyploids may also be related to which genes are expressed at a given time or tissue. This is likely relevant when comparing the assembled gene space for diploid and polyploid transcriptomes, as we do here. Our non-model reference transcriptomes are built from the expressed genes in each sample rather than comparison to a reference genome collection. Thus, only genes and alleles that are expressed will be captured in our assemblies and observed in our comparisons. Not all genes or alleles in a polyploid need to be expressed at one time and the overall diversity of the transcriptome at any given time may look more like a diploid, with other alleles being expressed at different times or tissues. Indeed, differential homoeolog silencing is well characterized in polyploid plants (Adams et al., 2003; Coate and Doyle, 2010) and may reduce the sampled transcript diversity of a polyploid genome. If this is the case, we would expect that sampling across more tissues, development times, and environments would lead to greater sampling of the polyploid gene space. Although RNA spike-ins and cell counting may improve differential expression analyses (Visger et al., 2019), capturing the full genome diversity of non-model polyploid species from RNA-seq assemblies remains an additional challenge.

Our pilot study of RNA-seq sampling of diverse species in the field demonstrated some familiar and unique challenges. Building on our past experience with extracting RNA from diverse species (Barker et al., 2008, 2016; Dempewolf et al., 2010; Der et al., 2011; Lai et al., 2012; Dlugosch et al., 2013; Hodgins et al., 2014; Mandáková et al., 2017; Qi et al., 2017; Yang et al., 2017; An et al., 2019; Carpenter et al., 2019), we developed an approach for this study to obtain high quality RNA from field samples. We found that flash freezing leaves in liquid nitrogen *in situ* for later RNA extraction worked well for our diverse samples. A few samples, especially *Polygonum cilinode*, yielded lower quality RNAwhich could potentially be related to leaf age at the time of sampling. Different RNA extractions methods will be needed to deal with the secondary compounds that are present in mature and senescing tissues. Recovering high quality RNA across a range of time points, in the field, from leaves of different ages will be a challenge for future studies.

Other challenges that will need to be overcome are associated with sampling at NEON sites. Sampling within NEON permanent plots is generally not allowed for collections outside of NEON’s own standard protocol, and therefore our sampling was limited to sites adjacent to NEON plots. This limitation raises some significant issues for researchers that wish to leverage data being collected within NEON sites. First, many NEON sites are located in areas where there is no similar adjacent field site available for sampling, due to land restrictions or ecological variation. We ultimately selected Harvard Forest because we could sample at sites other than NEON the plot itself. The second major issue is that sampling outside of the NEON plot means that there is no guarantee of continued access to plant populations in the future. There is a great opportunity for ecologists and evolutionary biologists to leverage the wealth of data that NEON is generating for our community. However, access for researchers that wish to conduct RNA and DNA sampling of plants (and other organisms) within NEON sites is an essential issue that requires further development across the network. Sequencing costs will continue to decline over the planned 30 year life span of NEON, and strategies to accommodate sequencing for plants and other eukaryotes will offer opportunities to greatly expand large scale studies at the intersection of ecology and evolution.

## ACKNOWLEDGEMENTS

We thank Manisha V. Patel at Harvard Forest for sampling logistics, George M. Ferguson at the University of Arizona Herbarium (ARIZ) for help with identifications and voucher deposits, and Jason Steel at the Biodesign Institute at Arizona State University for RNA-seq preparation and sequencing. This project was supported by NSF #1550838 to M.S.B. and K.M.D and NSF #1750280 to K.M.D.

## AUTHOR CONTRIBUTIONS

H.E.M., K.M.D., and M.S.B. conceived and designed the experiments. H.E.M. and S.A.J collected the samples, extracted the RNA, and collected the vouchers. H.E.M., E.W., Z.L., K.M.D., and M.S.B. analyzed the data. H.E.M., Z.L., K.M.D., and M.S.B. drafted the manuscript.

## DATA AVAILABILITY

Raw reads for all samples for 24 species are deposited on the NCBI Sequence Read Archive (SRP127805: https://www.ncbi.nlm.nih.gov/sra/SRP127805; BioProject: PRJNA422719). Assembled transcriptomes for each species are archived on Zenodo and available at https://doi.org/10.5281/zenodo.3727312.

